# New tools to study the interaction between integrins and latent TGFβ1

**DOI:** 10.1101/2023.01.26.525682

**Authors:** Michael Bachmann, Jérémy Kessler, Elisa Burri, Bernhard Wehrle-Haller

## Abstract

Transforming growth factor beta (TGFβ) 1 regulates cell differentiation and proliferation in different physiological settings, but is also involved in fibrotic progression and protects tumors from the immune system. Integrin αVβ6 has been shown to activate latent TGFβ1 by applying mechanical forces onto the latency-associated peptide (LAP). While the extracellular binding between αVβ6 and LAP1 is well characterized, less is known about the cytoplasmic adaptations that enable αVβ6 to apply such forces. Here, we generated new tools to facilitate the analysis of this interaction. We combined the integrin-binding part of LAP1 with a GFP and the Fc chain of human IgG. This chimeric protein, sLAP1, revealed a mechanical rearrangement of immobilized sLAP1 by αVβ6 integrin. This unique interaction was not observed between sLAP1 and other integrins. We also analyzed αVβ6 integrin binding to LAP2 and LAP3 by creating respective sLAPs. Compared to sLAP1, integrin αVβ6 showed less binding to sLAP3 and no rearrangement. These observations indicate differences in the binding of αVβ6 to LAP1 and LAP3 that have not been appreciated so far. Finally, αVβ6-sLAP1 interaction was maintained even at strongly reduced cellular contractility, highlighting the special mechanical connection between αVβ6 integrin and latent TGFβ1.

## Introduction

Transforming growth factor beta (TGFβ) 1 is an intensively studied cytokine of the TGFβ family with important functions in regulating the immune system and vascularization (Shull *et al*., 1992; Dickson *et al*., 1995). At the same time, excessive TGFβ1 signaling is associated with fibrotic tissue stiffening, for which therapeutic options are still limited (Kim, Sheppard and Chapman, 2018; Akhurst, 2017). TGFβs, in contrast to other members of the TGFβ family, are secreted in a latent state and require activation before they can bind their receptors (Robertson and Rifkin, 2016). The latent TGFβ complex consists of a dimeric TGFβ cytokine and a dimeric prodomain, called latency associated peptide (LAP). LAP 1 contains an RGD peptide that allows binding of cell adhesion receptors of the integrin family. Replacing this RGD peptide with RGE in knock-in mice prevents integrin binding to LAP1 and causes a phenotype similar to a *Tgfb1* knockout (Yang *et al*., 2007). This demonstrates, at the same time, the need for TGFβ1 activation and the relevance of integrin-LAP1 binding for this activation. One of the most established mechanisms for integrin-dependent TGFβ1 activation is mechanical pull via αVβ6 integrin (Munger *et al*., 1999; Dong *et al*., 2017). Such mechanical activation is potentially supported by the high affinity between LAP1 and αVβ6 integrin that allows to withstand the mechanical pull on the ligand-integrin-actin axis (Dong *et al*., 2014). While the ligand-integrin interface on this axis is well characterized on structural and biochemical level, much less is known about adaptations of the cytoplasmic integrin β6 tail. Special adaptations of the β6 tail compared to the tails of other integrins might be needed in order to transmit forces from the actin cytoskeleton onto LAP1 in order to enable TGFβ1 activation.

Integrin tails are short compared to their extracellular part (human β6: around 40 vs. 690) and recruit a plethora of adapters that are necessary for integrins to fulfill their functions (Bachmann *et al*., 2019). For integrin activation, binding to both talin and kindlin has emerged as a crucial factor (Pinon *et al*., 2014; Theodosiou *et al*., 2016; Fischer *et al*., 2021; Orré *et al*., 2021). Studies in keratinocytes have shown that the β6 tail only recruits kindlin1, but not kindlin2 (Bandyopadhyay *et al*., 2012). At the same time, binding of a β6-peptide to surface immobilized talin1 showed stronger binding compared to β3-peptide binding to talin1 (Kukkurainen *et al*., 2014). These observations support the possibility of particular adaptations of αVβ6 compared to other integrins. However, the relevance of such adaptations for LAP1 binding and TGFβ1 activation remains to be tested. Given the complex interactions between integrin adapters and their vast numbers, it might be particularly important to perform such experiments in a cellular context. Studying the interaction between latent TGFβ1 and αVβ6 integrin in cell biological experiments is complicated due to the structural organization of the LAP1/TGFβ1 complex. Labeling at the N-terminus interferes with the extracellular anchorage of the latent complex and the C-terminus is buried within the TGFβ cytokine. However, the TGFβ1 cytokine is not necessarily needed to study the interaction between LAP1 and integrins. Based on structural analysis, we identified a shorter part of LAP1 that contains the integrin-binding RGD peptide of LAP1 and grafted it onto a chimeric construct of a GFP and the Fc-region of human IgG. Additionally, we identified the analogous sequences of LAP2 and LAP3 and created such short LAP-GFP-IgG (sLAP) constructs for all LAPs. These tools allowed us to study the interaction of different integrins with all sLAPs. We demonstrate a unique interaction between αVβ6 and sLAP1, based on a mechanical rearrangement of immobilized sLAP1 by αVβ6 integrin. Additional experiments emphasized the need of kindlin, but not talin, for sLAP1 binding and mechanical perturbations revealed a remarkable robustness of the αVβ6-sLAP1 interaction.

## Results

### Developing new tools for studying integrin-LAP interaction

In order to study latent TGFβ – integrin interaction in more detail, we decided to develop new tools that facilitate this analysis. Structural analysis of available latent TGFβ1 structures (PDB: 3RJR, 5FFO, 6GFF) showed that a part of LAP1 (amino acids 105-263) is not in direct contact with the TGFβ1 cytokine. Instead, this part contains the RGD peptide and the cysteines involved in dimerization of latent TGFβs. Moreover, this region of LAP1 consists of a series of secondary structures that form a coordinated, stabilized globular fold that promises to fold without need of additional chaperones. We decided to use this reduced, smaller sequence of LAP1 and to combine it with a construct combining a GFP, the Fc region of human IgG, and a signaling peptide for secretion (Fig. 1A). Our lab established similar constructs in the past to study the binding between the tyrosine kinase c-Kit and its ligand (Tabone-Eglinger *et al*., 2014). Additionally, based on sequence alignment and structural modeling (SFig. 1B), we identified the equivalent parts of LAP2 and LAP3 and created sLAP2 and sLAP3 constructs (sequence as shown in SFig. 1A). Next, we transfected cells with the different sLAP constructs and harvested the supernatant. Western blotting of this supernatant showed the dimerized chimeric proteins of the small LAPs combined with GFP-IgG (sLAP) at the expected molecular weight (Fig. 1B). Moreover, structural prediction of sLAP1 and comparison to a crystal structure of latent TGFβ1 (PDB: 6GFF) supported the potential of sLAP1 to mimic the structural arrangement of latent TGFβ1 (root mean square deviation (RMSD) of atomic position: 0.701 Angstroms; Fig. 1C). Next, we used sLAP1 for coating glass coverslips for cell biological experiments. To ensure specific substrate coating we coated coverslips with protein A and passivated remaining, uncoated areas with albumin. Finally, these substrates were covered with sLAP1 that is recruited via the high-affinity interaction between the coated protein A and IgG in sLAP1. This procedure will also ensure that sLAP1 is immobilized on the substrate in a reproducible topological orientation, allowing integrin-sLAP1 binding. Next, we used NIH 3T3 fibroblasts and transfected them to express different integrins of the αV family that were discussed to activate TGFβ1 (Robertson and Rifkin, 2016). These fibroblasts express low levels of αVβ3 integrin (Bachmann *et al*., 2020) and showed no cell spreading and low cell attachment on sLAP1 coated substrates (SFig. 1C), ensuring that binding and spreading of transfected cells is due to the exogenously expressed integrin and not due to endogenous cell adhesion receptors. Culturing these transfected cells on sLAP1 coated coverslips and analysis with TIRF microscopy showed a lack of αVβ1-clustering on sLAP1, while αVβ3, αVβ5, and αVβ6 integrin all showed clustering (Fig. 1D). Integrin αVβ5 expressing cells showed a lack of cell spreading compared to cells expressing αVβ3 and αVβ6. This is probably explained by the special set of integrin adapters recruited by αVβ5 integrin (Baschieri *et al*., 2018; Lock *et al*., 2018; Zuidema *et al*., 2018), specifically by the lack of paxillin recruitment (Soto-Ribeiro *et al*., 2019). Quantitative analysis of the area of clustered integrins confirmed the impression that αVβ3, αVβ5, and αVβ6 are able to bind to sLAP1 and to cluster on this substrate while αVβ1 integrin under these experimental conditions was less able interact with sLAP1 (Fig. 1E). While this clustering analysis indicated a similar potential of αVβ3, αVβ5, and αVβ6 integrin for sLAP1 binding, analysis of the sLAP1 coating revealed a remarkable difference. The sLAP1 coating was frequently remodeled underneath αVβ6 expressing cells, but not under cells expressing other integrins. The spatial appearance of the remodeled sLAP1 suggested a mechanical pulling of cells on the substrate. Such a mechanical explanation for the sLAP1 rearrangement is reminiscent of the model of αVβ6-mediated TGFβ1 activation by mechanical forces on the LAP/TGFβ1 - αVβ6 integrin - actin axis (Dong *et al*., 2017; Munger *et al*., 1999).

**Fig. 01:**
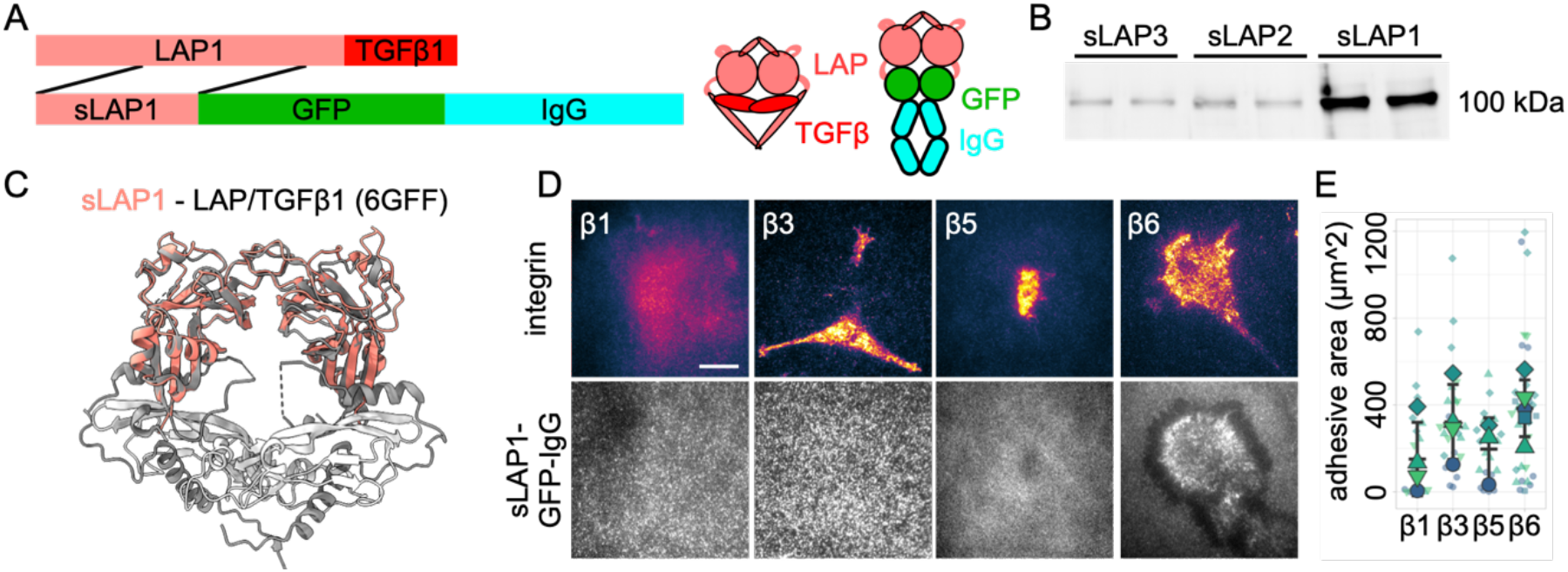
Chimeric sLAP-GFP-IgG as a tool to study integrin-LAP interactions. (A) Visualization of the organization of full-length TGFβ1 and of sLAP1-GFP-IgG proteins. (B) Non-reducing western blot of secreted chimeric sLAP-GFP-IgG proteins from cell culture medium. Two biological replicates per LAP are shown. (C) Structural alignment of a predicted Alphafold2 model of sLAP1 (shown in salmon) with protein structure of latent TGFβ1 (PDB: 6GFF; LAP1 in gray, cytokine in white) shows high overlap (RMSD: 0.701 angstroms). (D) NIH 3T3 fibroblasts were transfected with indicated plasmids (β1: β1-Scarlet, β3: β3-Scarlet, β5: β5-mCherry, β6: β6-Scarlet) and cultured for 2 hrs on glass coverslips, coated with sLAP1. Fixed cells were imaged with TIRF microscopy. Scale bar: 10 μm. (E) Analysis of area of clustered integrins for experiments performed as shown in (D). Small symbols stand for single measurements, big symbols for mean average from independent experiments. Middle bar is mean average from all experiments, upper and lower bar are standard deviation.

### Integrin αVβ6 rearranges exclusively sLAP1 coating

The consistent rearrangement of sLAP1 by αVβ6, but not by other integrins, motivated us to analyze the interaction of αVβ6 integrin with sLAPs in more detail. Specifically, we wondered if αVβ6 rearranges also other immobilized sLAPs. LAP1 and LAP3 share an RGD-Lxx(I/L) peptide with an unusual high affinity for αVβ6 integrin, while the SGD peptide in LAP2 does not favor strong αVβ6 binding ((Dong *et al*., 2014); see also SFig. 1A). The same RGD/SGD-peptides are present in our chimeric sLAP proteins. The interaction of integrins with these RGD/SGD peptides is often studied with short, synthesized peptides. In contrast, our sLAP proteins present the RGD/SGD peptides in their normal structural environment, similar to full-length LAPs. We used sLAP1-3 and coated glass substrates as we did before and cultured αVβ6 integrin expressing cells on these substrates (Fig. 2A). Cells showed reduced adhesion and spreading on sLAP2 compared to sLAP1 and sLAP3, indicating the relevance of the RGD-αVβ6 interaction for these processes. At the same time, only sLAP1 was rearranged by αVβ6 integrin (Fig. 2A, B). This difference between sLAP1 and sLAP3 was rather surprising given their similar integrin-binding RGD peptides. However, experiments with sLAP1-RGELATI and sLAP1-RGDGATG confirmed the necessity of the established high-affinity RGD-Lxx(I/L) peptide for substrate rearrangement of sLAP1 by αVβ6 integrin (SFig. 2B, C). However, the lack of sLAP3 rearrangement implies that this peptide is not sufficient for causing αVβ6-mediated rearrangement. Potentially, regions up- or downstream of the RGD-Lxx(I/L) peptide that differ between LAP1 and LAP3 are responsible for these different phenotypes (see discussion).

**Fig. 02:**
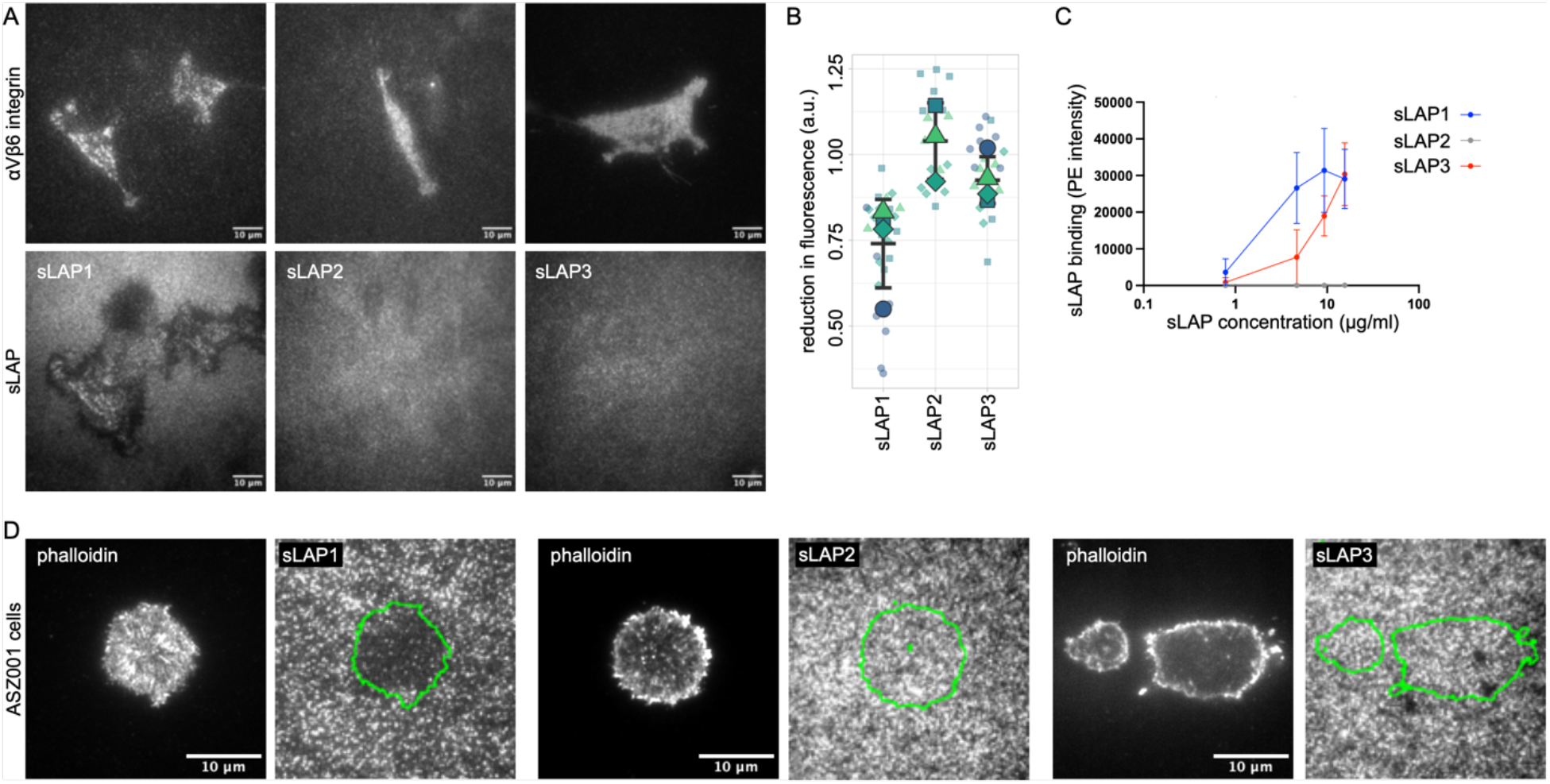
Integrin αVβ6 rearranges immobilized sLAP1, but not sLAP2 or sLAP3. (A) NIH3T3 cells expressing β6-Scarlet were cultured on substrates coated with indicated sLAPs and imaged with TIRF microscopy. Please note the rearranged sLAP1 coating underneath the cells, but the lack of rearrangement for sLAP2 and sLAP3. (B) Quantification of loss in fluorescence underneath cells compared to the intact substrate coating. Small symbols stand for single measurements, big symbols for mean average from independent experiments. Middle bar is mean average from all experiments, upper and lower bar are standard deviation. (C) Binding of β6-GFP expressing NIH3T3 cells to different concentrations of indicated sLAPs was quantified with flow cytometry (0.78, 4.67, 9.33, and 15.56 μg/ml). Average from 3 independent experiments with bars indicating standard deviation. (D) ASZ001 cells expressing αVβ6 integrin endogenously were cultured on coverslips coated with indicated sLAPs and imaged with TIRF microscopy (actin stained with phalloidin Alexa 647). Outline of cells are shown with green lines in images of sLAPs coating. Please note the removed sLAP1 underneath the cell in contrast to sLAP2 and sLAP3 coated conditions.

To characterize differences between binding of sLAPs to αVβ6 integrin in more detail, we performed a flow-cytometry based ligand binding experiment with αVβ6-expressing NIH3T3 cells, incubated with soluble sLAPs at different concentrations. Detection of surface-bound sLAPs with phycoerythrin (PE) labeled antibodies showed no binding of sLAP2 at any concentration, while sLAP1 and sLAP3 both bound strongly to αVβ6 expressing cells (Fig. 2c). However, sLAP1 showed higher binding capacity to αVβ6 integrin, potentially reflecting a higher affinity for αVβ6 integrin compared to sLAP3. Finally, we wanted to confirm that the rearrangement of sLAP1 is not limited to cells overexpressing exogenous αVβ6 integrin. Therefore, we used the ASZ001 murine cell line, a model for invasive basal cell carcinoma (Kuonen *et al*., 2018) that expresses αVβ6 integrin endogenously (SFig. 2A), and cultured them on substrates coated with the different sLAPs. Here again, we observed a consistent rearrangement of sLAP1, but not of either sLAP2 or sLAP3 (Fig. 2D).

### Dynamic and regulation of αVβ6-sLAP1 binding

Given the unique rearrangement of immobilized sLAP1 by αVβ6 integrin and the analogy of this process to the mechanical activation mechanism of TGFβ1 by αVβ6 integrin, we set out to analyze this process in more detail. We cultured NIH3T3 fibroblasts expressing β6-Scarlet on sLAP1 coated substrates and imaged them with live-cell TIRF microscopy starting 2 hrs after cell seeding (Fig. 3A). Similar to fixed samples, sLAP1 substrates underneath cells were already extensively rearranged at this time point. During the observation period, αVβ6 integrin clustered on areas of intact sLAP1 substrate coating and started to rearrange these regions. Compared to classical focal adhesions, αVβ6 was more dispersed in the membrane and formed clusters that were more irregular (Fig. 3A and SFig. 3A). At the same time, we observed that αVβ6 integrin clustering did not necessarily lead to sLAP1 rearrangement (see for example t = 45 min, sLAP1 compared to cell outline). This lack of correlation between active, clustered αVβ6 integrin and sLAP1 rearrangement could imply a potential regulation of αVβ6 integrin downstream of integrin activation. As a first test for this hypothesis, we transfected NIH3T3 cells with mutants of β6-integrin that affect either talin or kindlin binding. Mutating Tyr to Ala in the NxxY motifs of cytoplasmic β-integrin tails (see also Fig. 3B and SFig. 3C) is an established method to reduce binding of either talin (proximal NPxY) or kindlin (distal NxxY) (Saltel *et al*., 2009; Moser *et al*., 2008). At the same time, mutating Tyr to Phe in the proximal NPxY motif preserves aromatic interaction between the β-integrin tail and the talin F3-domain but prevents Tyr phosphorylation-induced binding of Dab2 (Yu *et al*., 2015). Dab2 binding to the phosphorylated integrin tail competes with talin and leads to inactivation and recycling of the integrin. Accordingly, we transfected cells with β6-Y762A to reduce talin binding, β6-Y762F to improve talin binding, and β6-Y774A to reduce kindlin binding and tested the ability of these mutants to bind sLAP1 in a flow cytometry-based ligand binding assay. After normalizing the amount of cell-bound sLAP1 to the surface levels of the respective αVβ6 integrin mutants, we observed no differences in sLAP1 binding for mutants affecting talin binding (Y762A and Y762F), but a consistent reduction for Y774A (Fig. 3C). Thus, kindlin but not talin seems to regulate binding of sLAP1 in this sLAP1 binding assay.

**Fig. 03:**
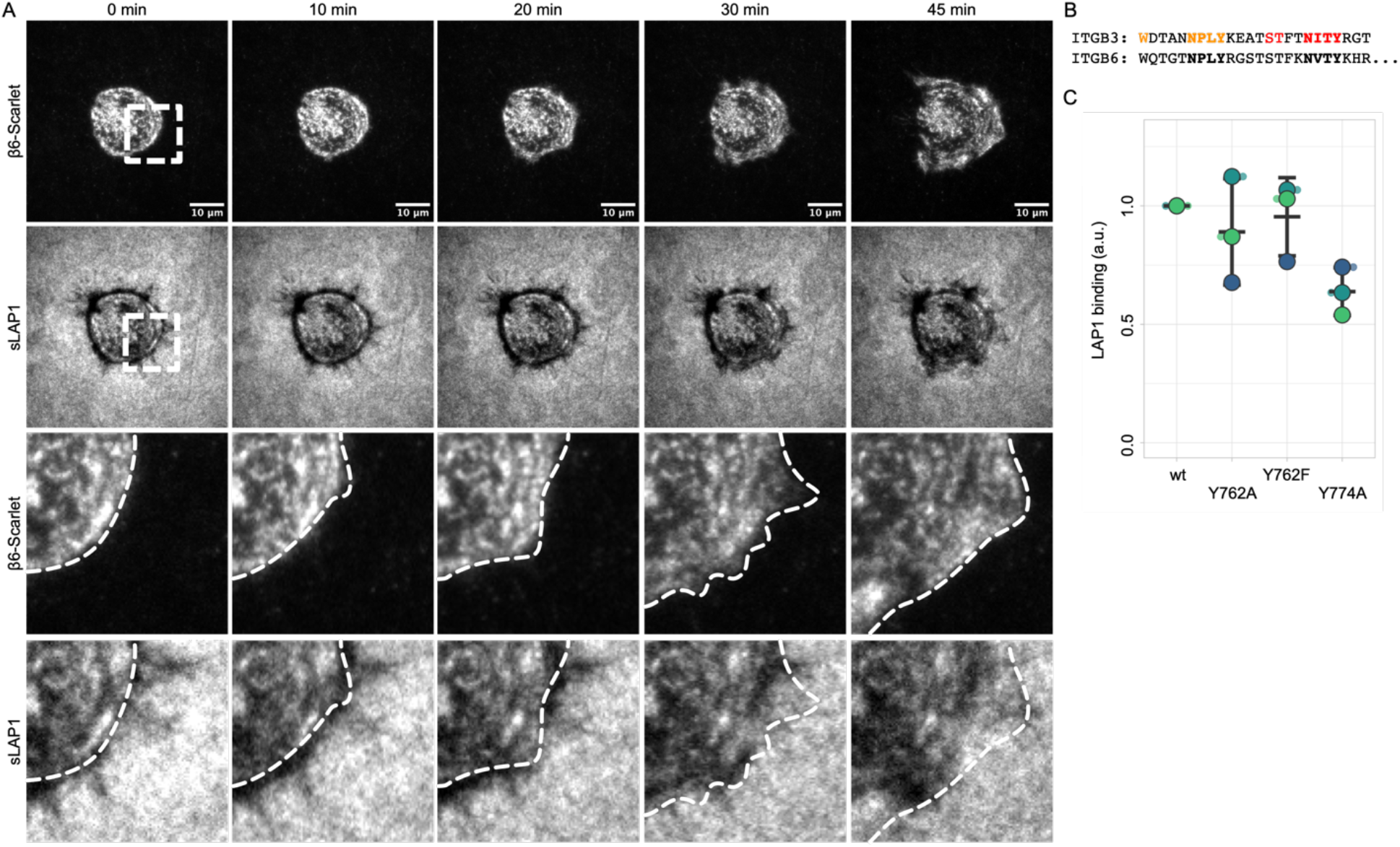
Dynamic regulation of αVβ6-sLAP1 rearrangement. (A) NIH3T3 cells expressing β6-Scarlet were cultured on sLAP1 coated substrates and imaged with TIRF microscopy starting 2 hrs after cell seeding (1 image / 5 minutes). White boxes in overview images indicate zoom-in regions shown in the lower rows. White dashed lines in zoom-in views indicate cell outline. Please note the clustered αVβ6 integrin at 45 min in the cell periphery without rearrangement of the underlying sLAP1 substrate. (B) Sequence alignment of cytoplasmic tail of β3- and β6-integrin. NxxY motifs are highlighted in bold, important talin-binding residues in orange and kindlin-binding residues in red (see also SFig. 3A). (C) NIH3T3 cells were transfected with mutants of β6-integrin affecting talin (β6-Y762A/F) or kindlin binding (Y774A) and incubated with sLAP1. Cell surface-bound sLAP1 was labeled with PE and measured with flow cytometry. Fluorescent intensity was normalized to β6 surface levels and β6-wt condition. Small symbols stand for single measurements, big symbols for mean average from independent experiments. Middle bar is mean average from all experiments, upper and lower bar are standard deviation.

Based on these experiments, it appears that kindlin and/or downstream targets of kindlin are regulators of sLAP1 binding. Future experiments with talin and kindlin knockout (KO) cells will establish if this finding also applies to the rearrangement of immobilized sLAP1.

### sLAP1-αVβ6 integrin binding is not affected by mechanical perturbations

The mechanical activation model of TGFβ1 via αVβ6 integrin implies strong mechanical pull on the LAP1-αVβ6 axis. The rearrangement of sLAP1 by αVβ6 seems reminiscent of the mechanical pull involved in this TGFβ1 activation model. Therefore, we tested the relevance of actin contractility for sLAP1 rearrangement. We cultured β6-Scarlet expressing NIH3T3 cells on sLAP1 coated substrates, either in presence of the myosin inhibitor blebbistatin or the ROCK pathway inhibitor Y27632. After 2 hrs, we fixed the cells and visualized the actin cytoskeleton. Both drugs had a strong effect on cell shape and the actin cytoskeleton compared to the untreated control (Fig. 4A). The untreated control also highlighted a strong rearrangement of sLAP1 in αVβ6 integrin expressing cells as observed before in figure 1–3. This rearrangement was characterized by a removal of sLAP1 from the substrate visible by the loss of GFP fluorescence underneath and next to the cells. Such a removal of sLAP1 was not easily detectable for blebbistatin and Y27632 treated cells. At the same time, cells treated with these inhibitors showed a strong accumulation of sLAP1 underneath the cells that was not observed to such a degree for untreated cells. This could imply that αVβ6 integrin is able to bind and to accumulate sLAP1 even when contractility is reduced, but that clearance of sLAP1 from the cell surface (for example by endocytosis and/or integrin inactivation and subsequent ligand unbinding) is perturbed due to the contractility inhibition. This hypothesis will be tested in more detail in future.

**Fig. 04:**
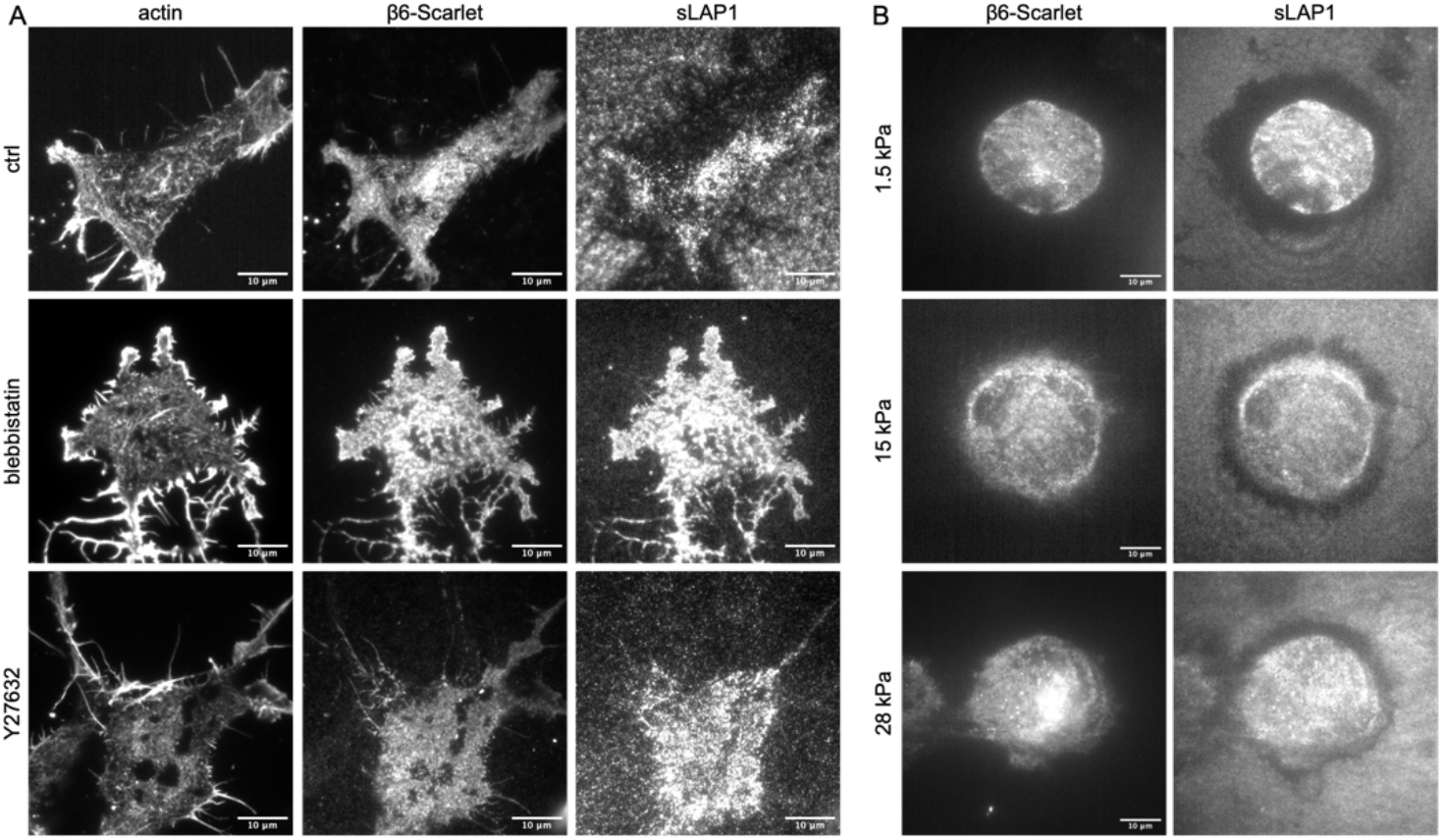
sLAP1 accumulation beneath cells is independent of actin contractility and substrate stiffness. (A) β6-Scarlet transfected NIH3T3 cells were cultured on sLAP1 coated substrates in presence of 50 μM of blebbistatin or of Y27632 where indicated. After fixation, actin cytoskeleton was visualized with phalloidin Alexa Fluor 647. (B) Elastic substrates (polydimethylsiloxane, PDMS) with indicated Young modulus were coated with sLAP1 and NIH3T3 cells expressing β6-Scarlet were cultured on them for 4 hrs before fixation and imaging with widefield fluorescence microscopy. Please note the similarities in cell spreading and rearrangement irrespective of the substrate stiffness.

To further characterize the interaction of αVβ6 integrin and sLAP1 under conditions with reduced cell contractility and less mechanical forces on ligand-integrin bonds, we cultured αVβ6 expressing cells on elastic substrates coated with sLAP1. We used substrates with stiffnesses at 1.5 kPa, 15 kPa, and 28 kPa to cover a wide range of physiological stiffnesses compared to our experiments performed before on unphysiologically stiff glass substrates. On all substrates, we observed a strong rearrangement of sLAP1, irrespective of the substrate stiffness. This result was unexpected given that substrates with a stiffness below 5 kPa typically cause a strong reduction in cell spreading and adhesion (Elosegui-Artola *et al*., 2016; Bachmann *et al*., 2020). At the same time, these results go in line with the ability of αVβ6 to bind and accumulate sLAP1 even when actin contractility is strongly reduced (Fig. 4A). And finally, αVβ6 is needed to activate TGFβ1 at physiological stiffnesses at 1.5 kPa and below. The ability of αVβ6 to bind and rearrange sLAP1 at 1.5 kPa as observed here could reflect this ability of αVβ6 to apply strong mechanical pull on ligands even in soft tissue environments.

## Discussion

We introduced new tools for studying the interaction between integrins and LAPs. These sLAPs proteins are easy to produce in cell culture and are versatile tools for different cell biological experiments. Using these tools, we made a number of unexpected observations with important consequences for TGFβ activation and signaling.

### Regulation of αVβ6-sLAP1 binding

There are reasons to argue that αVβ6 mainly evolved to activate TGFβ1 and that this is a major physiological function of this integrin. The affinity of αVβ6 integrin is around 200x higher for a LAP1-RGD peptide compared to a fibronectin-RGD peptide (Dong *et al*., 2014). Moreover, KO mice for *Itgb6* and for *Tgfb1* share an increased tendency for inflammation and a protection from fibrosis (Meecham and Marshall, 2020). However, fully mimicking the Tgfb1 KO mice requires the simultaneous KO and/or inhibition of αVβ6 and αVβ8 integrin (Aluwihare *et al*., 2009). The special nature of the LAP1-RGDLATI peptide with the additional binding of Leu and Iso into hydrophobic pockets in β6 integrin is potentially a structural/biochemical example for a possible co-evolution of αVβ6 integrin and the LAP/TGFβ complex. However, whether the cytoplasmic β6 tail also evolved special adaptations to allow TGFβ1 activation remains to be tested. Using sLAP1 as a new tool, we started to address this question. We could show a unique rearrangement of immobilized sLAP1 via αVβ6 integrin that can serve as a readout for mechanical interactions between this ligand and integrin. For now, we could confirm the relevance of both RGD and LATI binding to αVβ6 integrin for rearrangement of immobilized sLAP1 (SFig. 2B, C). Live-cell imaging showed the rearrangement of immobilized sLAP1 in the cell periphery. However, not every αVβ6 cluster led to sLAP1 rearrangement, possibly indicating that this ability of αVβ6 is regulated downstream of integrin activation and clustering. We tested β6 mutants affecting talin or kindlin binding and observed a need for kindlin binding (based on β6-Y774A mutation) to ensure sLAP1 binding. In future, we will use talin and kindlin KO cells to confirm this observation and to test the relevance of these adapters for the rearrangement of immobilized sLAP1. In addition, we will analyze downstream targets of talin and/or kindlin for their involvement in sLAP1 rearrangement, specifically paxillin, FAK, and vinculin. Paxillin has been shown to be recruited to integrins via kindlin (Theodosiou *et al*., 2016) and talin (Pinon *et al*., 2014), and talin-dependent paxillin recruitment is necessary to stabilize integrin adhesions (Ripamonti *et al*., 2021). At the same time, paxillin interacts closely with FAK and their interaction regulates integrin dynamics (Bachmann *et al*., 2022; Zaidel-Bar *et al*., 2007; Ballestrem *et al*., 2006). Vinculin supports force transmission between the integrin-talin-actin axis (Elosegui-Artola *et al*., 2016; Humphries *et al*., 2007; Thievessen *et al*., 2013) and could be expected to facilitate αVβ6-mediated mechanical TGFβ1 activation. The relevance of these adapters will be tested for sLAP1 rearrangement, as well as for TGFβ1 activation by using TGFβ reporter cell lines (Tesseur *et al*., 2006).

Another indication for a special mechanical LAP1-αVβ6 interaction stems from our observation that sLAP1 rearrangement is not affected by substrates stiffness. Experiments using different ligands and integrins have shown that a substrate stiffnesses below 5 kPa cause a drastic decrease in cell spreading and adhesion force (Elosegui-Artola *et al*., 2016; Bachmann *et al*., 2020), even when studying αVβ6-expressing cells on fibronectin (Elosegui-Artola *et al*., 2014). In contrast, we observed cell spreading and strong sLAP1 rearrangement at 1.5 kPa, again emphasizing the special nature of the LAP1-αVβ6 interaction. Studying the relevance of β6 adapters as mentioned above will also shine light on the mechanism allowing αVβ6 to withstand such mechanical perturbations.

### αVβ6 rearrangement of sLAP1 but not sLAP3

LAP1 and LAP3 share similar RGD peptides that bind with high affinity to αVβ6 integrin (Dong *et al*., 2014) and αVβ8 integrin (Ozawa *et al*., 2016). While the RGD peptide is also found in other ligands, LAP1 and LAP3 have an RGD that is followed by a Lxx(L/I) sequence (LAP1: LATI, LAP3: LGRL). The Leu/Iso residues bind into hydrophobic pockets of the respective β subunit and support the high affinity interaction (Dong *et al*., 2014). Based on these structural and biochemical insights, and supported by experimental data (Annes, Rifkin and Munger, 2002), it is widely assumed that TGFβ3 is activated similarly to TGFβ1. Therefore, we were quite surprised to observe differences between αVβ6-sLAP1 and αVβ6-sLAP3 interaction. These differences included the mechanical rearrangement of immobilized sLAP1 but not sLAP3, and a reduced binding capacity of soluble sLAP3 compared to sLAP1 in a flow cytometric ligand binding assay. Interestingly, some experiments demonstrating similarities of αVβ6 binding to either the LAP1-RGD or the LAP3-RGD sequence used short peptides that comprised not much more but the RGD-Lxx(L/I) sequence (Dong *et al*., 2014). In contrast, our sLAP constructs present the respective RGD peptides integrated into the bigger LAP complex similar to the full-length protein. Interestingly, the sequence surrounding LAP1-RGD is rather hydrophobic, while LAP3-RGD is surrounded by charged, hydrophilic residues (see also SFig. 1A). These different environments up- and downstream of the respective RGD peptides could modulate their binding to integrins, in particular when mechanical forces are applied to the integrin-RGD bond. In this regard, it might be worth noting that the specificity-determining loop 2 (SDL2) of β6 integrin contacts LAP1 and is rather hydrophobic (Dong *et al*., 2017). This hydrophobic signature of the β6 SDL2 might interfere with binding to the more hydrophilic nature of the LAP3-RGD peptide and explain the reduced binding and lack of αVβ6-mediated sLAP3 rearrangement. At the same time, the hydrophilic environment of the LAP3-RGD peptide might facilitate binding to other integrins with a more hydrophilic SDL2.

### Integrin-sLAP1 binding beyond αVβ6

Integrin αVβ6 and αVβ8 are established as activators of TGFβ1 in both cell culture assays using luciferase reporter cell lines (Dong *et al*., 2014), as well as in mouse experiments (Aluwihare *et al*., 2009). These studies imply that other integrins have no role in activating TGFβ1. At the same time, there is compelling data in mouse models for different fibrotic diseases that TGFβ1 activation by other integrins is crucial for TGFβ1 activation and subsequent fibrotic progression (Reed *et al*., 2015; Henderson *et al*., 2013; Noskovicova *et al*., 2021). Here, we observed a unique ability of αVβ6 integrin to rearrange immobilized sLAP1. However, while no other integrin was able to achieve such a rearrangement, both αVβ3 and αVβ5 showed prominent clustering on immobilized sLAP1. This observation raises the possibility that these integrins might also be able to activate TGFβ1 under certain conditions, in particular in fibrotic tissue. Fibrotic tissue is characterized by ECM stiffening, but also an increased activity of ECM-degrading proteases (Menou, Duitman and Crestani, 2018). Proteolytic cleavage of LAP1 leading to TGFβ1 release is the established mode for αVβ8 integrin mediated TGFβ1 activation (Mu *et al*., 2002). Increased proteolytic activity in a fibrotic environment might make this mechanism of TGFβ activation more accessible for other integrins. In parallel, ECM stiffening and increased cellular contractility could modulate the LAP1-binding capacity of integrins like αVβ3 and αVβ5 and, thereby, increase their ability for TGFβ1 activation. We showed recently that the ligand binding capacity of αVβ3 integrin is increased by cellular contractility and mechanical pull on the integrin-ligand bond (Bachmann *et al*., 2020). Force-induced LAP1 binding by integrins like αVβ1, αVβ3, and αVβ5 could explain the conflicting data about their ability for TGFβ1 activation mentioned above. It is noteworthy that these integrins have not been observed to activate TGFβ1 in experiments using low-contractile HEK cells (Dong *et al*., 2014) or in mice without fibrotic challenge (Aluwihare *et al*., 2009) while data for their relevance for TGFβ1 activation stems from fibrotic settings (Reed *et al*., 2015; Henderson *et al*., 2013). Testing the ability of cells of different contractility to interact with sLAP1 might offer a way to test this hypothesis.

## Methods

### Cell culture, constructs, and transfection

NIH3T3 cells were cultured in Dulbecco’s Modified Eagle Medium (DMEM, Pan-Biotech, Germany) supplemented with 10% fetal calf serum (FCS, HyClone, USA), glutamine, and 1% penicillin/streptomycine at 10% CO2 and 37°C. The NIH3T3 clone expressing low amounts of endogenous αVβ3 integrin was described recently (Pinon *et al*., 2014). ASZ001 cells were a gift from François Kuonen (University of Lausanne, Switzerland). ASZ001 cells were cultured in 154CF medium (ThermoFisher), supplemented with 2% chelated FCS (Chelex 100 molecular biology grade resin, BioRad, USA), 1% penicillin/streptomycin, and 0.060 mM CaCl_2_. Transfections were carried out with JetPEI (Polyplus Transfection) according to the manual. The plasmid for expressing β5-mCherry was a gift from Guillaume Montagnac and Francesco Baschieri (Gustave Roussy, France, (Baschieri *et al*., 2018)). Scarlet-I was obtained via PCR from pmScarlet-i_C1 deposited by Dorus Gadella (addgene #85044, (Bindels *et al*., 2017)). β1-Scarlet and β3-Scarlet were created by replacing GFP from the SapMar-β1-GFP construct described previously (Soto-Ribeiro *et al*., 2019; Egervari *et al*., 2016) and the pcDNA3-β3-GFP construct described previously (Ballestrem *et al*., 2001). A SapMar-β6-Scarlet plasmid was created by replacing β1 in the SapMar-β1-Scarlet plasmid by β6 obtained by a PCR from the integrin beta6 EGFP-N3 plasmid deposited by Dean Sheppard (addgene #13593) and a SapMar-β6-GFP plasmid was created analogously by replacing β1 in SapMar-β1-GFP. For creating chimeric sLAP proteins, we introduced a NotI and a XhoI site between the signaling peptide and the GFP of GFP-IgG constructs that were described previously (Tabone-Eglinger *et al*., 2014). LAP domains for creating sLAP1-3 were obtained from plasmids for the respective full-length human TGFβs, deposited by Gavin Wright (TGFΒ1, addgene #52185, (Sun *et al*., 2015)), F. Xavier Gomis-Ruth (TGFΒ2, addgene #128500, (Marino-Puertas *et al*., 2019)), and Charles Gersbach (TGFΒ3, addgene #52580, (Brunger *et al*., 2014)). The shorter, reduced LAP fragments for creating sLAPs (bold letters in SFig. 1A) were obtained via PCR from the TGFβ constructs mentioned above and were inserted via NotI-XhoI digest into the NotI-XhoI-GFP-IgG construct created before. Mutations of sLAP1 and of β6 in SapMar-β6-GFP were created by site-directed mutagenesis and digestion and ligation using unique restriction sites. The sequences of all constructs were verified by sequencing.

### Antibodies and reagents

Inhibition experiments were performed with blebbistatin (Sigma-Aldrich, USA) or with the ROCK inhibitor Y27632 (Sigma-Aldrich, USA) at concentrations as indicated. Cells were fixed with 2% paraformaldehyde (Sigma-Aldrich, USA) in PBS. Actin cytoskeleton was visualized with phalloidin-Alexa Fluor 647 (ThermoFisher, USA). For flow cytometry, integrin β6 was stained as described previously with an antibody developed by the antibody facility at the University of Geneva (Kessler *et al*., 2022). Integrin αV was stained with a commercial antibody (ThermoFisher, USA, clone RMV-7, 1:1000 diluted). Labeling of primary antibodies was visualized with anti-mouse PE and anti-rat PE (both Southern Biotech, USA, 1:1000 diluted). Actin in living cells was visualized with SiR actin (Spirochrome, Switzerland).

### Producing chimeric sLAP proteins, coating substrates, and cell seeding

HEK-293T cells (obtained from ATCC, #CRL-3216) were transfected with the calcium-phosphate method according to established protocols. Specifically, 1.5×10^6 cells were seeded in 10 cm dishes in DMEM + 10% FCS. The next day, 10 μg DNA was combined with 125 μl of 2 M CaCl_2_ and filled up with ddH_2_0 to a volume of 1 ml. This volume was slowly combined with 1 ml of 2x Hepes Buffered Saline pH 7.0 (ThermoFisher, USA) on a shaker to ensure rapid mixture of the solutions. After 15 min incubation, the DNA solution was added to the cultured HEK cells, together with 25 μl of 10 mM hydroxychloroquine-sulfat (SigmaAldrich, USA). The following day, cells were washed with PBS, followed by incubation in Opti-MEM Reduced Serum Medium (ThermoFisher, USA). After 5 days, medium was collected and centrifuged to pellet floating cells. Supernatant was filtrated for sterilization. The presence of the sLAP proteins was verified by western blotting according to standard protocols and by staining with anti-human HRP antibodies (Southern Biotech, USA). Concentration of sLAP proteins was quantified by dot blotting and using purified human Fc-IgG (SigmaAldrich, USA) as concentration standard.

For coating of substrates with sLAP proteins, glass coverslips were cleaned with ethanol and incubated with 20 μg/ml of protein A (BioVision, USA) for 1 hr at room temperature or 4°C over night. After washing, coverslips were incubated with 0.5% w/v of human serum albumin (SigmaAldrich, USA) for 30 min at room temperature, followed by another washing step. Afterwards, coverslips were incubated for 30 min at room temperate with supernatant of the respective sLAP proteins. Coverslips were washed a final time and used for experiments.

For cell seeding on these substrates, cells were detached with trypsin/EDTA solution (ThermoFisher, USA) and collected by centrifugation. Cells were washed twice with PBS to remove trypsin/EDTA as well as FCS. Afterwards, cells were seeded onto sLAP coated substrates in DMEM without serum and cultured for 2 hrs before fixation.

Elastic substrates (ibidi, Germany) were coated as described above and cell seeding was also performed as described with the exception that cells were cultured for 4 hrs.

### Flow cytometry

For detection of surface levels of integrins, 1×10^5^ cells were kept in solution and on ice and were incubated with the respective antibodies for 30 min. Cells were pelleted and washed and incubated with secondary antibodies again for 30 min on ice. Antibodies were diluted in DMEM supplemented with 1% w/v of bovine serum albumin (AppliChem, Germany). After the secondary antibody incubation, cells were again pelleted and washed and were analyzed with an AttuneNxT flow cytometer (ThermoFisher, USA). Cells were gated based on forward and side scatter to detect living cells and for singlet cells. For experiments with transfected cells, cells were additionally gated based on GFP expression to analyze only transfected cells. Ligand binding assays were performed in the same way but the incubation with the primary antibody was replaced by incubation with the respective sLAP supernatant at the indicated concentrations; incubation was performed at 37°C for 30 min. Surface-bound sLAP was detected with anti-human PE antibodies that bound directly to the Fc-IgG part of the chimeric sLAP proteins (SouthernBiotech, USA). PE signals were analyzed with FlowJo software (BD Biosciences, USA). For Fig. 3C, sLAP binding and αVβ6 integrin surface levels were measured for the same condition. sLAP binding was normalized by the amount of surface expressed αVβ6 for the different conditions. Finally, all conditions were normalized to the level of sLAP binding in β6-GFP wt transfected condition.

### Microscopy and image analysis

All imaging was performed on a Nikon Eclipse Ti with perfect focus system, 100x/1.49 NA oil objective and an Andor-EMCDD iXon897 camera. Imaging was either performed with TIRF illumination or in widefield mode. Live-cell imaging was performed in a heated chamber at 37°C and 5% CO2 and with cells cultured in glass-bottom dishes. Cells were cultured in DMEM + 10% FCS for 2 hr before imaging. For imaging, medium was changed to F12 buffer (ThermoFisher, USA) with 10% FCS. Images were prepared and analyzed using the Fiji software package (Schindelin *et al*., 2015) and custom-written macro codes (available from https://github.com/Mitchzw).

### Statistics and software

Plots were prepared with SuperPlotsOfData (Goedhart, 2021) or with Prism (GraphPad, USA) in case of Fig. 2C. Error bars always indicate standard deviation based on averages from at least 3 independent experiments.

Structural models of sLAP1-3 were created with Alphafold2 and ColabFold (Jumper *et al*., 2021; Mirdita *et al*., 2022). Visualization of structural models and protein structures was done in ChimeraX (Pettersen *et al*., 2021; Goddard *et al*., 2018).

## Acknowledgements

We are grateful to Monica Julio Barreto for help with molecular cloning. We thank the Bioimaging Core Facility, the Flow Cytometry Facility, and the Antibody Facility of the University of Geneva for their support and help. We acknowledge UCSF ChimeraX, developed by the Resource for Biocomputing, Visualization, and Informatics at the University of California, San Francisco, with support from National Institutes of Health (R01-GM129325) and the Office of Cyber Infrastructure and Computational Biology, National Institute of Allergy and Infectious Diseases.

## Data availability

All data that is part of this manuscript will be made available before final publication via Yareta, the archive repository of the University of Geneva.

## Supplementary Material

**Supplementary Figure 01:**
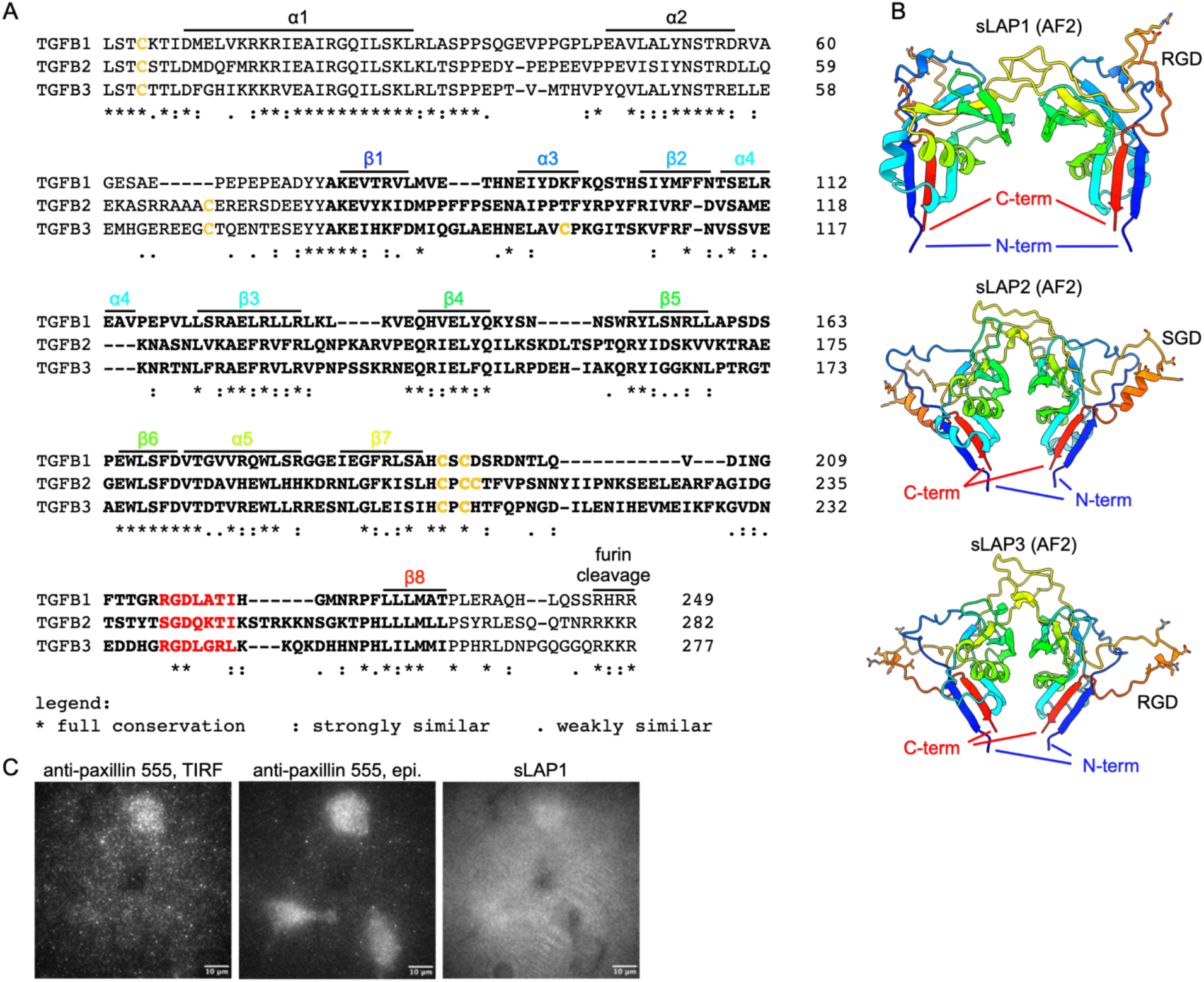
(A) Sequence alignment of LAP parts of human TGFβ1-3. Cysteines are highlighted in yellow, regions used for sLAPs in bold, and RGD/SGD peptides in red. Numbering of secondary structures is based on 6GFF (Liénart *et al*., 2018); their color coding is according to sLAP1 shown in (B). (B) Alphafold2 predictions of sLAP1, sLAP2, and sLAP3. Secondary structures are rainbow colored with N-terminus in blue and C-terminus in red. Please note the dimerization of the LAP proteins via cysteines following β7 (“bowtie region”), the presence of the RGD/SGD peptides in a flexible loop, and the availability of both N- and C-terminus for adding molecular tags. (C) Mock-transfected NIH3T3 cells were cultured on sLAP1 coated substrates and were stained against endogenous paxillin. Please note the presence of cells in the field of view as indicated by the epifluorescent image of paxillin (middle panel), while these cells neither rearrange sLAP1 nor establish prominent focal adhesions.

**Supplementary Figure 02:**
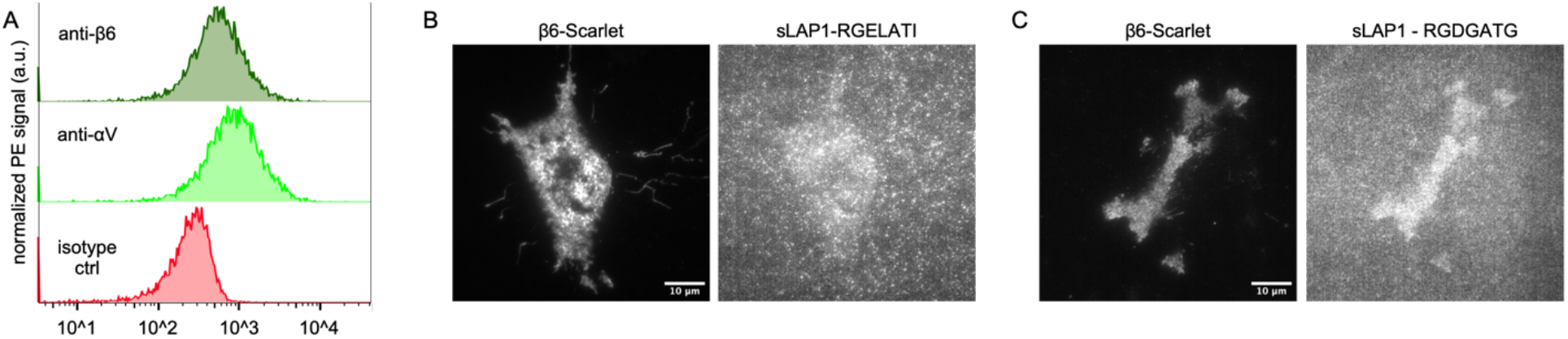
(A) Flow cytometric analysis of surface expression of αV integrins and of β6 integrin. (B, C) β6-Scarlet expressing NIH3T3 cells cultured on sLAP1 substrates where the high-affinity RGD-LATI peptide was mutated to (B) RGE-LATI or (C) RGD-GATG.

**Supplementary Figure 03:**
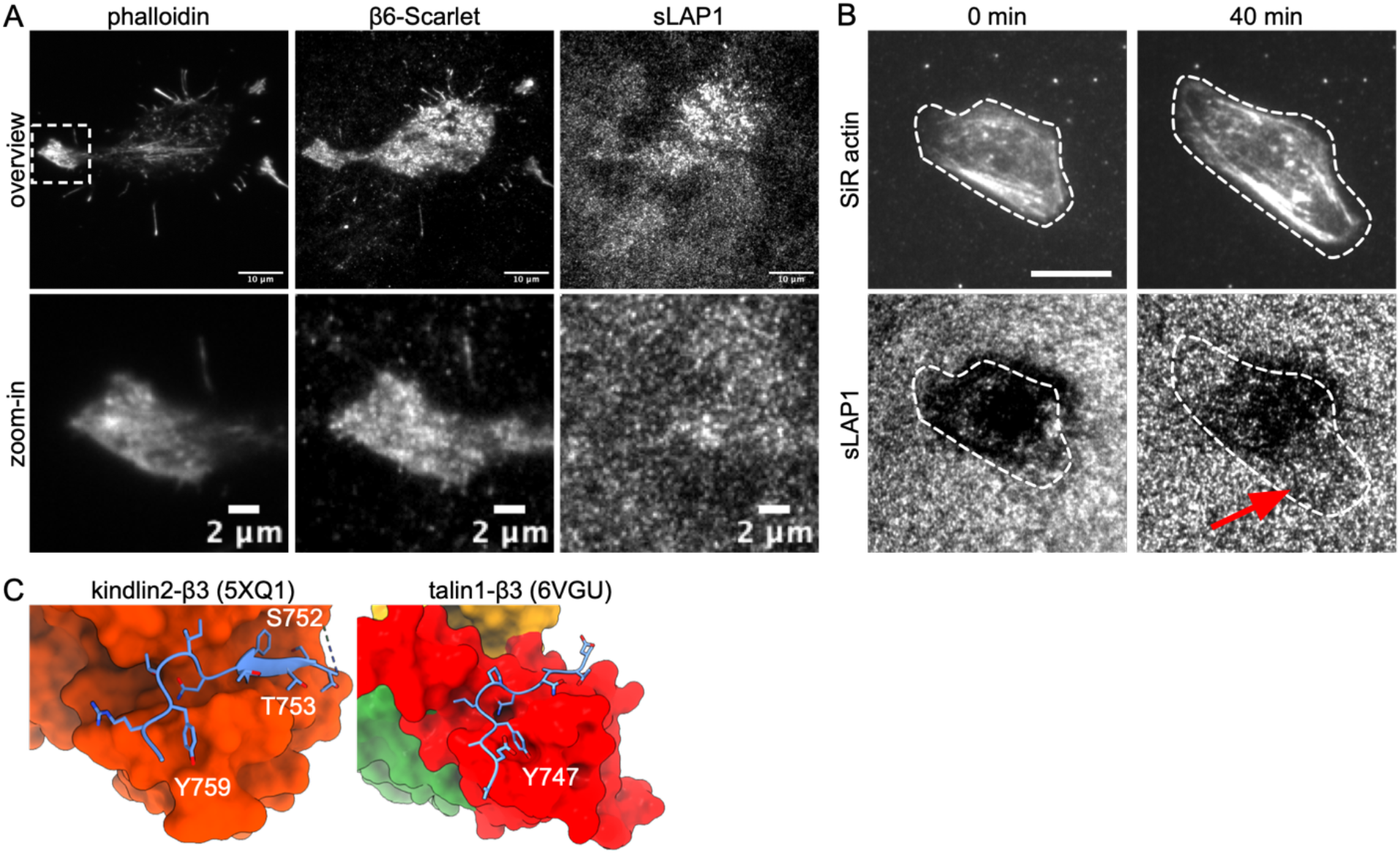
(A) β6-Scarlet expressing NIH3T3 cells were cultured on sLAP1 coated substrate, fixed, and stained for actin with phalloidin 647. White box is shown as zoom-in in the lower row. (B) ASZ001 cells were incubated with 1 μM of SiR actin to visualize the actin cytoskeleton in living cells. Cells were imaged for 45 min every 2.5 min. Outline of cell at respective time points is shown as white dashed line. Please note the reduced sLAP1 fluorescence in areas occupied by the cell during the time lapse imaging (red arrow). Scale bar: 10 μm. (C) Structural analysis of integrin β3 tail (bright blue) binding to F3-domains of kindlin2 (PDB: 5XQ1, orange) or of talin1 (PDB: 6VGU, red; F2 in green, F1 in yellow). Please note the binding of the Tyr of the proximal and distal NxxY motifs to respective pockets in talin and kindlin. Also highlighted are S752 and T753 that are involved in forming a β-sheet with kindlin (see also Fig. 3B) and that have been shown to be important for integrin binding to kindlin (Soto-Ribeiro *et al*., 2019).

